# CIB2 as a Ca^2+^ Sensor for Auditory Mechano-Electrical Transduction and Linked to Genetic-Heterogeneity of *TMC1* in Hearing Loss

**DOI:** 10.1101/2024.07.24.604958

**Authors:** Shaoxuan Wu, Lin Lin, Qiaoyu Hu, Xuebo Yao, Hongyang Wang, Shuang Liu, Qingling Liu, Yuehui Xi, Yuzhe Lin, Jianqiao Gong, Ruixing Hu, Wei Zhan, Yi Luo, Guang He, Zhijun Liu, Wei Xiong, Qiuju Wang, Zhigang Xu, Fang Bai, Qing Lu

## Abstract

Non-syndromic sensorineural hearing loss is characterized by genetic heterogeneity, leading to potential clinical misdiagnosis. *TMC1*, a unique causative gene associated with deafness, exhibits variants with autosomal dominant and recessive inheritance patterns. *TMC1* codes for the transmembrane channel-like 1 (TMC1), a key component of the mechano-electrical transduction (MET) machinery for hearing. However, the molecular mechanism of Ca^2+^ regulation in MET, which is essential for sound perception, remains unclear. CIB2, another MET component associated with deafness, can bind with calcium and may play a role in MET gating. Our study reveals that the TMC1-CIB2 complex undergoes a Ca^2+^-induced conformational change, highlighting the crucial role of Ca^2+^ in MET. We identified a vertebrate-specific binding site on TMC1, named CR3, that interacts with *apo* CIB2, with the binding interface linked to hearing loss. Using an in-vivo animal model, we demonstrated that disruption of the calcium-sensing site of CIB2 perturbs the MET channel conductivity, confirming its role as a Ca^2+^ sensor for MET. Additionally, after systematically analyzing the hearing-loss variants, we observed that dominant mutations of TMC1 are often located near the ion pore or at the binding interfaces with the Ca^2+^ sensor CIB2. This provides a detailed mechanism of the genetic heterogeneity of *TMC1* in hearing loss at an atomic level.

## Introduction

Hearing loss affects an estimated 5% of the world’s population, making it a prevalent sensory deficit in humans. Over 150 genes and 8000 variants have been implicated in the pathogenesis of hearing loss^1^, highlighting the heterogeneity of this condition and the need for gene-specific or variant-specific phenotype studies. *TMC1* (OMIM*606706) is a commonly implicated gene in genetic hearing loss ^2–7^, with two distinct phenotypes. Variants in *TMC1* have been associated with both autosomal dominant and autosomal recessive hearing loss ^8–11^, located separately at DFNA36 and DFNB7/11, and present distinct clinical phenotypes. DFNB7/11 is characterized by congenital severe to profound hearing loss, with a minority of patients presenting with moderate hearing loss. On the other hand, DFNA36 is featured by acquired or postlingual progressive hearing loss ^9,11–13^. Accurate genetic variant classification is crucial for proper genetic diagnoses, but this remains a major challenge in the post-genome era.

In the auditory system, the mechano-electrical transduction (MET) protein machinery in the cochlear hair cells is responsible for sound perception^14–16^. The MET machinery is located in the actin-based protrusions, known as stereocilia, which are organized in a staircase-like structure. Soundwaves trigger the deflection of stereocilia, causing the MET channels to open, leading to the influx of K^+^ and Ca^2+^ ions and altering receptor potential ^17^. Several proteins have been identified as the components of the mammalian hair cell MET machinery, including the transmembrane-like channel 1 and 2 (TMC1 and TMC2) ^11,18–20^, cadherin 23 (CDH23) ^21^, protocadherin-15 (PCDH15) ^22,23^, lipoma HMGIC fusion partner-like 5 (LHFPL5) ^24,25^, transmembrane inner ear (TMIE) ^26,27^, and calcium and integrin-binding protein 2 and 3(CIB2 and CIB3) ^28–31^. TMC1 and TMC2 are pore-forming subunits of the MET channel and can act as a mechanosensor^19,20,32–34^. CIB2 and CIB3, as cytosolic proteins, play a crucial role in MET complex assembly^30,31,35^. Variants of *TMC1* and *CIB2* genes have been linked to hearing loss ^8,11,29,36–39^. Recent studies have determined the TMC1/2-CALM1-TMIE complex structure in worms^40,41^. However, the molecular mechanisms underlying mammalian MET assembly, maintenance, and gating kinetics for hearing remain to be fully elucidated.

The hair cell MET channel, in comparison to other mechanosensitive channels, exhibits remarkably fast open-close kinetics that enable it to respond to fluctuating sound input. The gating kinetics of the channel (activation, fast adaptation, and slow adaptation) are functionally modulated by biophysical and pharmacological factors including Ca^2+^. The roles of Ca^2+^ in MET gating are incredibly intriguing, including extracellular Ca^2+^ homeostasis and intracellular Ca^2+^-dependent adaptation ^42–46^. The concentration of Ca^2+^ in the scala media, at 20 μM, maintains the stereocilia’s stiffness and rigidity ^42,43,47^. An influx of Ca^2+^ modulates the activation and adaptation of the transducer current ^44–46^. CIB2, a Ca^2+^-binding protein and critical binding partner of TMC1, whose null mutation in mice result in abolished mechanotransduction currents, may play essential role of the intracellular Ca^2+^-mediated modulation on the MET channel kinetics.

CIB2 consists of four EF-hands (helix-loop-helix, Ca^2+^-binding motif) ^48^. Calmodulin and CaBP1, as well-studied members of the EF-hand-containing superfamily, are Ca^2+^ sensors for voltage-gated sodium channels and Ca^2+^ channels (CaV1.2 and CaV1.3, collectively called CaV) ^49–51^. After the opening of CaV channels, an increase of cytosolic Ca^2+^ levels lead to Ca^2+^-bound calmodulin modulating the conductivity of CaV and facilitating rapid Ca^2+^-dependent channel inactivation ^52^. Our study reveals that CIB2 is the Ca^2+^ sensor for TMC1 conformational changes and could be important for MET gating in fluctuating sound perception. Additionally, we have characterized disease mutations that interfere with the TMC1-CIB2 interactions through structural analysis, biochemical validation, and *in-vivo* assays, thoroughly investigating the clinical phenotypes from micro to macro.

## Results

### The complex structure of mammalian CIB2 and TMC1 cytosolic region 1 (CR1)

Previous studies have shown that two cytosolic regions (CRs) of TMC1 (CR1, amino acids 81– 130 & CR2, amino acids 303-347) (Fig. 1A) bind to CIB2/CIB3 and have reported the complex structures of TMC1 CR2-CIB3 ^29,30^. The recently reported complex structure of the worm TMC1-CALM1 complex lacks the details about the interaction between TMC1 CR1 and CIB2 due to resolution limitations ^40^. To gain a better understanding of the atomic level interaction between the mammalian TMC1 CR1 and CIB2, we co-crystallized the protein complex of mouse TMC1 CR1 and CIB2. We then successfully solved the mammalian complex structure at a resolution of 1.74 Å.

**Fig. 1.**
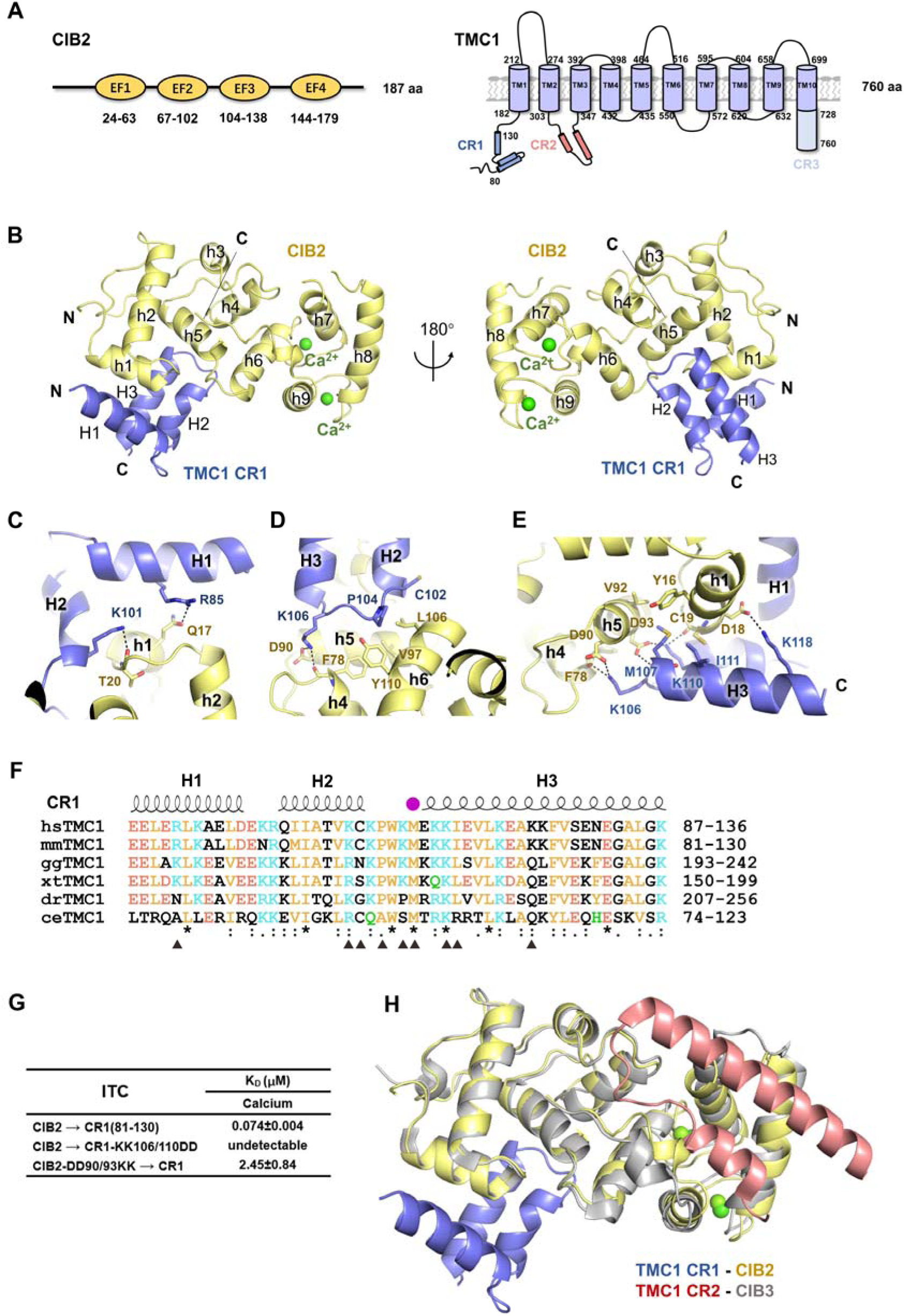
Crystal structure of the mammalian TMC1 CR1-CIB2 complex. (**A**) Schematic diagrams show the organization of the EF-Hand domains of CIB2, transmembrane helices, and cytosolic regions of TMC1. (**B**) Ribbon diagram representation of the TMC1 CR1-CIB2 structure as viewed from the front (left) and the back (right); secondary structure elements are labeled. (**C-E**) Detailed interactions between CIB2 and TMC1 CR1. Residues involved in protein interaction are shown with the stick model. CIB2 and TMC1 CR1 are colored in slate and pale yellow, respectively. Hydrogen bonds are shown as black dashed lines. (**F**) Sequence alignments of TMC1 CR1 from different species. The secondary structural elements of TMC1 CR1 determined from this work are shown above its alignment. Conserved residues are colored orange (nonpolar), green (polar), blue (positive charged), and red (negative charged). Specifically, residues indicated with black triangles are responsible for the interaction between CIB2 and TMC1 CR1; the dominant disease mutation site is indicated with a purple circle. hs: *Homo sapiens*; mm: *Mus musculus*; gg: *Gallus gallus*; xt: *Xenopus tropicalis*; dr: *Danio rerio*; ce: *Caenorhabditis elegans*. (**G**) Summary of the binding affinities between WT or structural mutants of CIB2 and TMC1 CR1. (**H**) Model alignment of TMC1 CR1-CIB2 with TMC1 CR2-CIB3 (PDB: 6WUD).

In the complex structure, the overall fold of CIB2 is very similar to other CIB family proteins, but the binding mode between CIB2 and TMC1 CR1 is entirely unexpected. CIB2 has nine α-helices (h1 to h9) forming four EF-hands, with only EF-hands 3 and 4 of CIB2 binding Ca^2+^. The CR1 of TMC1 folds into three short α-helices (H1 to H3) (Fig. 1B), binding with the N-terminal region of CIB2. In contrast, CR2 folds into two α-helices and contacts with a hydrophobic trench at the C-terminal of CIB3.

The interface between the TMC1 CR1 and CIB2 buries a surface area of about 740.9 Å^2^ between these two molecules. TMC1 CR1 contacts the N-terminal region of CIB2, involving h1, h2, and h4-h6. The TMC1 CR1-CIB2 interface is mainly mediated by electrostatic interactions, hydrogen bonds, and hydrophobic interactions. R85 and K101 of TMC1 form hydrogen bonds with Q17 and the main chain of T20 of CIB2, respectively (Fig. 1C). C102 and P104 from the linker between H1 and H2 of TMC1 contact with the hydrophobic groove formed by F78, V97, L106, and Y106 (Fig. 1D). In addition, K106, K110, and K118 of TMC1 form salt bridges with D90, D93, and D18 of CIB2, respectively (Fig. 1D). Hydrophobic interactions exist among M107, I111 from CR1 and Y16, C19, and V92 from CIB2 (Fig. 1E). The sequence alignment of the TMC1 CR1 region among different species indicates that the residues involved in TMC1 interaction are highly conserved in vertebrates but less conserved in worm TMC1 (Fig. 1F). Consistently, mutations (CIB2-DD90/93KK and CR1-KK106/110DD) that disrupt these polar interactions weakened or even abolished the TMC1 CR1-CIB2 interaction in isothermal titration calorimetry (ITC) assays (Fig. 1G). Based on the structure superimposition of TMC1 CR1-CIB2 and TMC1 CR2-CIB3 (Fig. 1H), we found that the two binding surfaces of CR1 and CR2 are located at opposite ends of CIB2/CIB3, suggesting the possibility of CR1 and CR2 binding to CIB2 simultaneously and the potential existence of additional undiscovered CIB2-binding sites. On the other hand, CIB2 belongs to the EF-hand family, which includes Calmodulin, a well-characterized calcium sensors that participates in various physiological processes ^53–55^. This led us to consider the calcium-binding property of CIB2’s EF-hands.

### Ca²□ mediated regulation on the binding between CIB2 and TMC1 cytosolic regions (CRs)

The high-resolution complex structure mentioned above prompted us to conduct an extensive biochemical characterization of CIB2’s binding to different TMC1 cytosolic regions and its interaction with calcium (Fig. 1A&2A&2B). Upon detailed examination of all cytosolic regions of TMC1, we discovered a new binding site in TMC1 C-terminal, named TMC1 CR3 (718-757, at the extreme C-terminal of TMC1 TM10). Sequence alignment showed that the CR3 region is only conserved among vertebrates, emerging with the presence of the inner ear (Fig. 2B). To quantitatively analyze the binding affinity, we performed ITC assays to measure the dissociation constant under conditions with or without calcium. In the presence of calcium, CIB2 bound to CR1 and CR2 with the dissociation constant (*K*_D_) values of ∼ 0.074 μM and ∼ 0.48 μM, respectively (Fig. 2C). Without calcium, the binding affinity of CIB2-CR1 and CIB2-CR2 decreased to ∼ 0.87 μM and ∼ 10.8 μM. Unlike CR1 and CR2, CR3 can only form complexes with CIB2 in the absence of calcium, with a binding affinity of ∼43.4 μM (Fig. 2C).

**Fig. 2.**
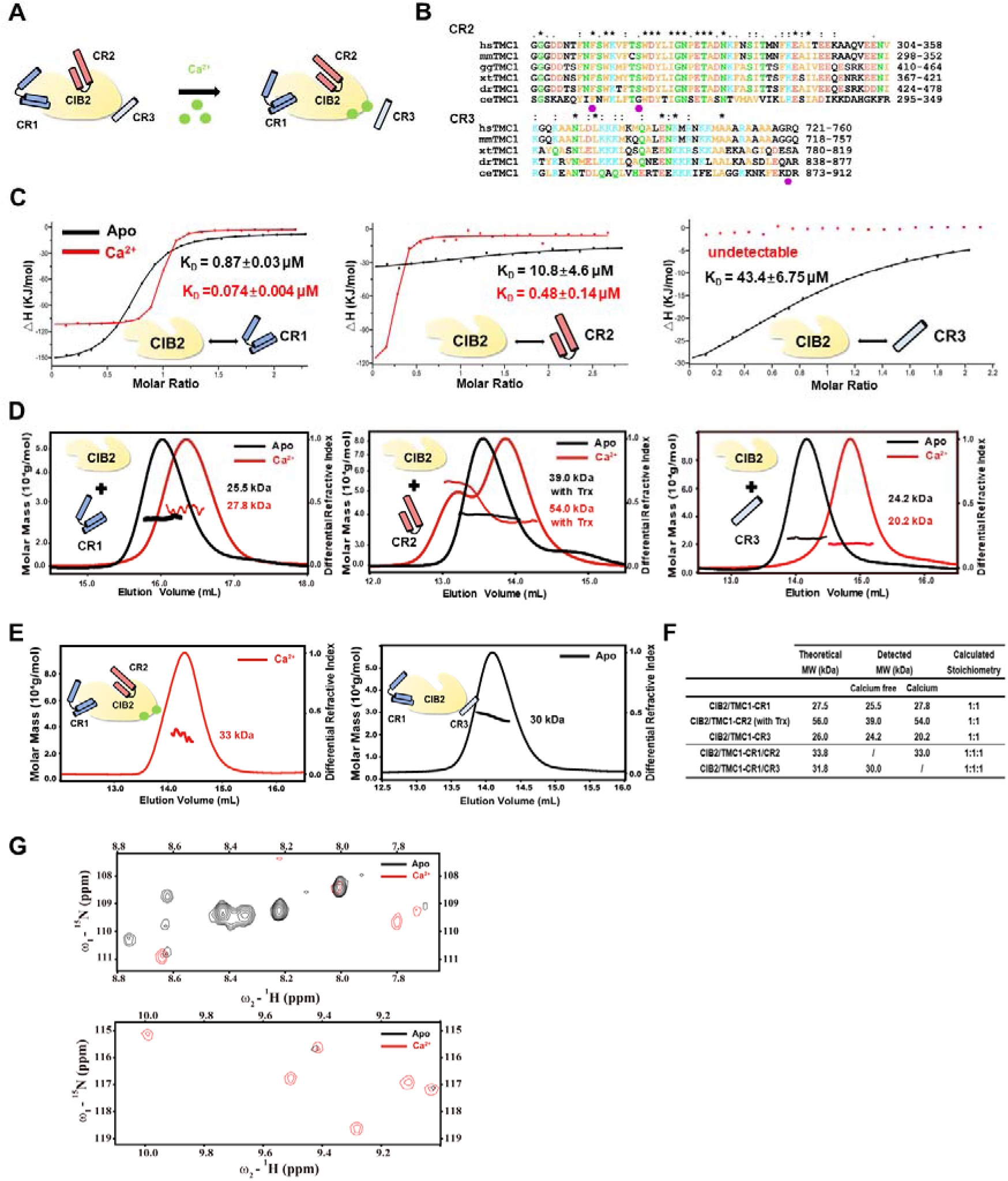
CIB2 interacts with CR1, CR2, and CR3 of TMC1, regulated by calcium. (**A**) Schematic diagrams show the interaction between CIB2 and three cytosolic regions of TMC1 and the conformational change of the complex after CIB2 binding Ca^2+^. (**B**) Sequence alignments of TMC1 CR2 and TMC1 CR3 from different species. Conserved residues are colored orange (nonpolar), green (polar), blue (positive charged), and red (negative charged). The dominant disease mutation sites are indicated with purple circles. hs: Homo sapiens; mm: Mus musculus; gg: Gallus gallus; xt: Xenopus tropicalis; dr: Danio rerio; ce: Caenorhabditis elegans. (C) ITC analysis shows that CIB2 binds to TMC1 in three cytosolic regions (CR) with different binding affinity in *apo* (black) and Ca^2+^ (red) states. 10 mM EDTA/CaCl_2_ were added in buffer. (**D**) SEC-MALS analysis of CIB2 binds to TMC1 CR1/CR2/CR3 in *apo* (black) and Ca^2+^ (red) states. Trx, thioredoxin. (**E**) SEC-MALS analysis of CIB2 with TMC1 CR1&TMC1 CR2 in Ca^2+^ state (red) and TMC1 CR1&TMC1 CR3 in *apo* state (black). (**F**) A summary of SEC-MALS results, showing the molecular weight and calculated stoichiometry. (**G**) An overlay plot of a selected region of 1H-15N HSQC spectrum of CIB2 and TMC1-CR1 complex. The *apo* state is showing in black, and Ca^2+^ state is showing in red. 10 mM EDTA/CaCl2 were added in buffer. ppm: parts per million.

To reveal the formation of the complex, we utilized size-exclusion chromatography coupled with multi-angle static light scattering (SEC-MALS) to determine the molar mass of the complex. First, upon introducing Ca^2+^ to the CIB2-CR1 complex, we observed a shift to a higher elution volume, suggesting the presence of Ca^2+^ induced a conformational change in the CIB2-CR1 complex (Fig. 2D). In the case of CIB2-CR2, we observed a molar mass increase and a ∼ 200-fold increase in binding affinity (Fig. 2C). These findings indicate a more significant conformational modulation on CIB2-CR2 mediated by Ca^2+^. Regarding the CIB2-CR3 complex, we found that CIB2 forms a complex with CR3 only in the absence of calcium, which is consistent with the ITC data. To understand the composition of the mixture of CIB2 with CR1 and CR2 under calcium conditions, we determined the molar mass and discovered a binding ratio of 1:1:1 for CIB2:CR1:CR2 (Fig. 2E&2F). This supports the prediction made by our structure analysis that both CR1 and CR2 can simultaneously bind to CIB2. Similarly, in the *apo* state (absence of calcium), the mixture of CIB2 with CR1 and CR3 also displayed a binding ratio of 1:1:1 for CIB2:CR1:CR3, indicating the simultaneous binding of CR1 and CR3 (Fig. 2E&2F).

To further validate the conformational changes induced by calcium in the CIB2-CR1 complex, NMR-based titration experiments were conducted (Fig. 2G). In the *apo* state, significant ^1^H-^15^N HSQC signals were observed at ω_2_-^1^H 8.0-8.8; these signals vanished upon Ca^2+^ binding to CIB2, demonstrating the exquisite sensitivity of NMR chemical shifts to Ca^2+^ binding. Meanwhile, in the Ca^2+^-bound state of the complex, residues show signals at ω_2_-^1^H 9.0-10.0, indicating conformational changes in the complex as a result of Ca^2+^ binding. To further consolidate the calcium-sensing ability of the TMC1-CIB2 complex, we assessed their preference for divalent ions. Our ITC experiments revealed that CIB2 has a binding affinity of ∼ 3.43 K_D_ for Ca^2+^, but no significant binding to Mg^2+^ (Fig S2A). The opposite conclusion compared to previous report^56^ may be attributed to the adoption of different experimental methods. Additionally, we identified the interaction of CIB2 and CR3 under Mg^2+^ state, which showed a comparable binding affinity to the *apo* state (Fig S2B). Inductively coupled plasma-Mass Spectrometry (ICP-MS) analysis showed that TMC1 CR1 and CIB2 complex binds to five times more Ca^2+^ than Mg^2+^ under cellular concentrations (10:1 ratio of Mg^2+^ to Ca^2+^, Fig S2C).

In summary, we identified a vertebrate-specific binding site on TMC1, named CR3, that interacts with *apo* CIB2. And Ca^2+^ regulates the binding between CIB2 and TMC1 cytosolic regions by modulating CR1 and CR2 interactions and dissociating CR3 (Fig. 2A). To understand how Ca^2+^ mediates MET channel function, we moved on to elucidate the calcium regulation on the complex of CIB2 and the full-length TMC1.

### The assembly of the mammalian full-length TMC1-CIB2 complex

Combing the complex structures of mammalian TMC1 CR1-CIB2 (Fig. 1B) and TMC1 CR2-CIB3 (PDB ID: 6WUD)^30^, biochemical characterization on the Ca^2+^-regulated association between TMC1 and CIB2 (Fig. 2), and the previous studies on TMC1 transmembrane regions that TMC1 assembles as a dimer and resembles TMEM16 ion channels^19^, we can generate the structural assembly model of mammalian TMC1 and CIB2 complex (Fig. 3A). Firstly, the interaction between CIB2 and full-length TMC1 was verified by the co-immunoprecipitation (Co-IP) experiment. We found the deletion of individual CR domains reduced the CIB2-TMC1 interaction in the absence of Ca^2+^ (Fig. S3A). However, even with both CR1 and CR2 deleted, CR3 still interacted with CIB2 in the absence of Ca^2+^, indicating that TMC1-CR3 is involved in the *apo* state of the complex (Fig. S3B).

**Fig. 3.**
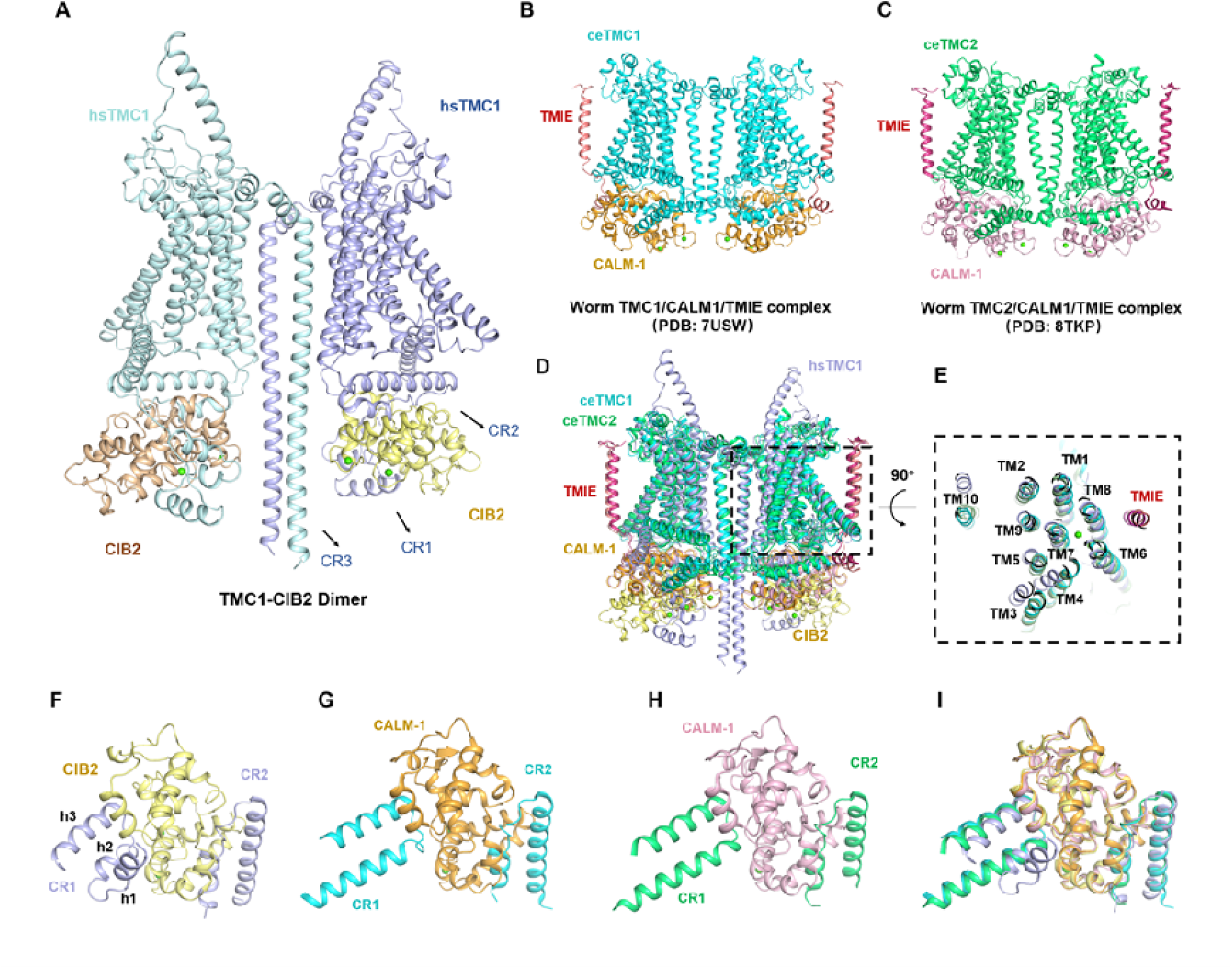
The assembly of the mammalian TMC1-CIB2 complex. (**A**) The structure model of TMC1-CIB2 dimer. The CR1, CR2, and CR3 segments of TMC1 and CIB2 are labeled. Th calcium ions are shown as green spheres. (**B**) The complex structure of *C.elegans* TMC-1/CALM-1/TMIE (PDB code: 7USW). (**C**) The complex structure of *C.elegans* TMC-2/CALM-1/TMIE (PDB code: 8TKP). (**D**) Superimposition of *C.elegans* TMC-1 complex, TMC-2 complex and human TMC1-CIB2 complex model. (**E**) The top-down view of the superimposition of the transmembrane regions of *C.elegans* TMC1, TMC2 complexes and human TMC1-CIB2 complex model. (**F-I**) Local structural comparison of the human TMC1 CR1/CR2 complexed with CIB2 (**F**), *C. elegans* TMC-1 cytoplasmic regions (CR1 and CR2) complexed with CALM1 (**G**) and *C. elegans* TMC-2 cytoplasmic regions (CR1 and CR2) complexed with CALM1 (**H**).

The structure model of the human TMC1 dimer was obtained using the deep-learning-based protein-protein interaction predictor AlphaFold-Multimer with default settings for model parameters and the selection of a complete genetic database ^57^. The predictions were ranked based on a model confidence metric, a weighted combination of predicted TM-score (pTM) and interface pTM. The whole structure of the TMC1-CIB2 dimer (Fig. 3A) was constructed by aligning the TMC1 and CIB2 onto the structure of CIB3-TMC1 CR2 (PDB ID: 6WUD) and substituting the TMC1 CR1 segment (amino acids 81-130) by the corresponding counterpart in our crystal structure of CIB2-CR1. The preparation of molecular dynamics (MD) input files of the TMC1-CIB2 dimer was conducted through the bilayer builder of CHARMM-GUI ^58^ (Fig. 3B). The protein complex with the membrane was employed as the starting point for the following all-atom MD simulations, which were performed using the GROMACS program^59^ and CHARMM36 force field ^60^.

As displayed in Fig. 3A, the structure of the TMC1-CIB2 dimer exhibits a symmetric conformation between two monomers, which are formed with the mediation of the TM10 and the CR3 helices (Fig. 3A). The CR1(at the N-terminal of TM1) and CR2 (between TM2 and TM3) of TMC1 embrace the CIB2 protein in the intracellular region, and the calcium ions bind to the EF-hand 3 and 4 in CIB2 (Fig. 3A). The cytoplasmic part provides sufficient space for the simultaneous binding of CR1 and CR2 with CIB2 in the calcium-bound state. The putative pore of TMC1, characterized in the previous study^19^, is formed by TM4-8.

We then performed a structural superimposition of the human TMC1-CIB2 complex model with the recently reported *C. elegans* TMC-1/CALM1/TMIE complex (Fig. 3B) and TMC-2/CALM1/TMIE complex structures^41^ (Fig. 3C). The overall dimer conformation and transmembrane region topology show similarity among the three complexes (Fig. 3D). All the transmembrane regions of ceTMC-1, ceTMC-2 and the hsTMC-1 protomer are composed of ten helices (Fig. 3E). However, significant variations are observed in the cytosolic regions, especially in CR1 and CR3, which are responsible for CIB2 association confirmed by our biochemical data. Our structure shows that the CR1 of human TMC1 consists of three short α-helices (h1 to h3), with h1 being absent in the worm TMC-1 and TMC-2 structures and sequences (Fig. 3F to 3I). On the other hand, the conformation of CR2 is conserved, folding into two α-helices and making contact with a hydrophobic trench at the C-terminal of CIB2/CALM (Fig. 3F to 3I). The other notable difference lies in the CR3 region, which forms a long helix extending from the TM10 helix in our complex structure model, but it is completely absent in both worm structures (Fig. 3E).

### The conformational change of full-length TMC1-CIB2 complex in response to Ca²□

To understand the molecular dynamic behavior of the full-length TMC1-CIB2 complex under the regulation by Ca²□, the algorithmic molecular dynamic-based simulation workflow was designed accordingly. The simulations were performed on the two conformational states of the TMC1-CIB2 complex, i.e., the *apo* state without Ca²□ and the Ca²□ bound *holo* state. For simulation of the Ca²□ state (Fig. S4A), the complex was equilibrated in a cubic box filled with TIP3P^61^ water molecules, some of which were replaced by sodium and chloride ions. Subsequently, energy minimization, NVT, and NPT equilibrations were conducted sequentially to stabilize the temperature and pressure of the system to 310 K and 1 bar, respectively. Finally, MD simulation was performed on the equilibrated structure for 300 ns under the NPT ensemble as in previous reports^62,63^. To simulate the *apo* state (Fig. S4B), we used two different strategies, changing the membrane tension and changing the membrane thickness, to mimic the mechanical signal. A more decisive design is to remove calcium ions to monitor the dynamic behavior based on our findings of calcium modulation. The following analyses were conducted on the most representative structure (center structure of a cluster) clustered from MD trajectories.

Detailed structural analysis revealed the following conformational change between the two states. First, in the MD-derived structure of TMC1-CIB2 Ca²^+^ form, the CR3 segment of TMC1 is away from CIB2, consistent with our biochemistry results of no detectable binding between CR3 and Ca²□ -CIB2. The Ca^2+^ binding involves the residues of D116, N118, D120, D127, D157, D159, D161, and D168 in CIB2 (Fig. S4A, left panel). In the *apo* state with membrane tension change, the CR3 region bends toward CIB2, thus forming multiple salt bridges (R759-D120, R759-D161, K746-D159, K742-D157, K742-D168, R744-E170, and R744-D171) with residues within and around the previous Ca^2+^-binding sites (Fig. S4B, left panel). Meanwhile, we also used another strategy to do *apo* state simulation with a thinner membrane used in simulating the opening process of MscL channel^64^. The electrostatic interactions formed between CR3 and CIB2 were also observed in the structure derived from the simulation (Fig. S4C, left panel). This provides the structural basis for the Ca^2+^-mediated modulation of the TMC1-CIB2 complex. Consistently, ITC results showed that although single mutations of Lys/Arg to Ala have little effect, triple substitutions of the positively-charged residues with Ala in CR3 significantly decrease its binding affinity with CIB2 (Fig. S4D&S7E).

### Disruption of the Ca²□ sensing element perturbs the channel conductivity in vivo

To consolidate our hypothesis, we conducted a series of mutagenesis experiments to assess whether the Ca²□ sensing element on CIB2 is essential for the proper function of MET. It is worth mentioning that MET was unaffected in *Cib2* knock-out vestibular hair cells (VHCs), and CIB3 could compensate for the loss of CIB2 in VHCs ^31^. Detailed structural analysis of the mammalian CIB2-TMC1 CR1 complex structure revealed that EF-hand 3 and EF-hand 4 of CIB2 each bind one Ca^2+^ in a canonical manner. In EF-hand 3, D116, N118, D120, F122, and D127 participate in Ca^2+^ coordination (Fig. 4A); in EF-hand 4, D157, D159, D161, K163, and D168 interact with the other calcium ion (Fig. 4B). It is assumed that substituting D116&D120 on EF-hand 3 and D157&D161 on EF-hand 4 with alanine should disrupt the Ca^2+^ coordination. We first verified the direct impairment of both double mutants in their binding with Ca^2+^, resulting in a ∼25-fold decrease for DD116/120AA and a ∼12-fold decrease for DD157/161AA compared to CIB2 WT (Fig. S5A). The ITC experiments further showed that both double mutants impair their binding with TMC1 CR1 in response to calcium (∼150-fold decrease for DD116/120AA in Fig. 2C&4A and ∼50-fold decrease for DD157/161AA in Fig. 2C&4B). These results suggested that the Ca^2+^ sensing element on CIB2 was essential for the interaction with TMC1.

**Fig. 4.**
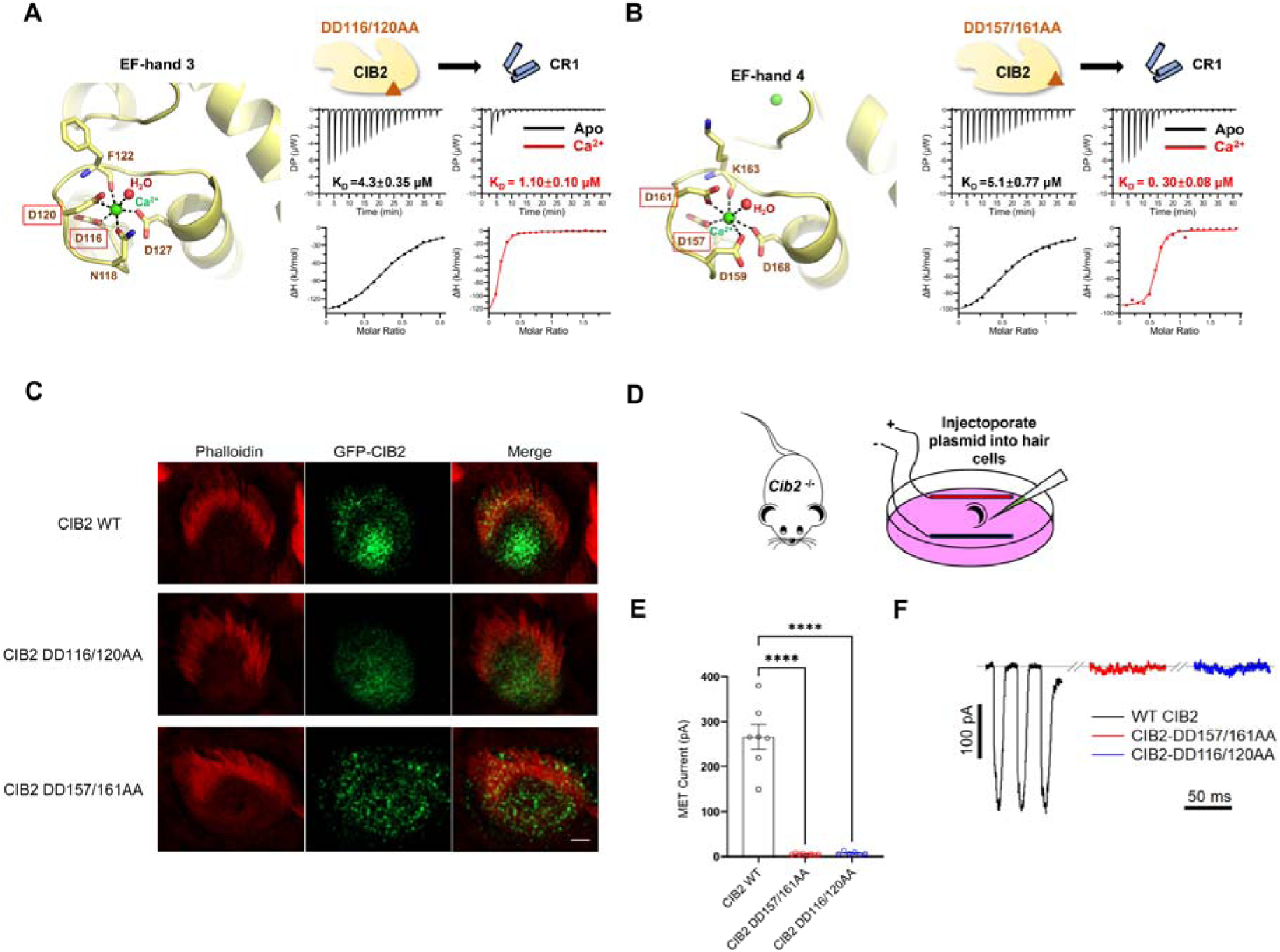
Calcium-sensing mutations disturb MET assembly and affect MET in mice OHCs. (**A, B**) The structural details of EF-hand 3 and EF-hand 4(left panel) and ITC results of double mutant residues of CIB2 interacting with CR1. The mutant residues are framed in red; th calcium ions are colored in green. ITC results are shown in apo state (black) and Ca^2+^ state (red), respectively. (**C**) Examples of OHCs from wild-type mice injectoporated with the indicated constructs and visualized for CIB2 (green) and phalloidin (red). Scale bar =1 μm. (**D**) A schematic drawing of injectoporation procedure to express genes in hair cells. (**E**) Quantification of the MET current amplitudes from recordings in P3 Cib2^-/-^ OHCs injectoporated with different CIB2 constructs as indicated. Shown are means ± SEM; ****, p < 0.0001. (**F**) Representative examples of MET currents recorded from OHCs in Cib2^-/-^ mice injectoporated with different CIB2 constructs as indicated.

We injectoporated wild-type CIB2 as well as two mutants, DD116/120AA and DD157/161AA, into wild-type mouse cochleae (Fig. 4C). Immunostaining results demonstrated that CIB2-DD157/161AA localized to the stereocilia, similar to CIB2-WT (Fig. 4C). In contrast, CIB2-DD116/120AA exhibited impaired protein trafficking and localized mainly in the cell body (Fig. 4C). Given the approximately 25-fold decrease in binding affinity of CIB2-DD116/120AA for Ca^2+^ (Fig. S5A), this result indicates that defective calcium-binding of EF-hand 3 led to protein mislocalization of CIB2.

We further performed MET current rescue experiments on *Cib2* knock-out mice to consolidate the function of Ca^2+^ sensing elements in vivo. We recorded the MET current in *Cib2*^-/-^ mice injectoporated with different CIB2 constructs (Fig 4D). CIB2-WT successfully restored the MET current in P3 *Cib2*-deficient outer hair cells (OHCs). In contrast, no MET current was detected in CIB2-DD116/120AA or CIB2-DD157/161AA overexpressing *Cib2*-deficient OHCs (Fig 4E), despite CIB2-DD157/161AA having proper stereocilia localization. Representative examples of MET currents recorded from OHCs are shown in Fig 4F. These in vivo experiments confirmed the significance of Ca^2+^ sensing elements on CIB2 and the MET channel by regulating the interaction between CIB2 and TMC1.

### The autosomal dominant hearing loss mutations of TMC1 cluster around the putative pore of the MET

Variants in *TMC1* have been associated with both autosomal recessive and autosomal dominant hearing loss (DFNA36 and DFNB7/11). More than 100 different DFNB7/11 variants and 8 DFNA36 variants have been identified (Table S3, combination of reported variants and families recruited in this study). This genetic heterogeneity poses a significant challenge for accurate diagnosis and effective treatment. However, our comprehensive characterization of the TMC1-CIB2 complex (Fig. 5A) has enabled us to match these different clinical phenotypes with the underlying pathogenesis at an atomic level.

**Fig. 5.**
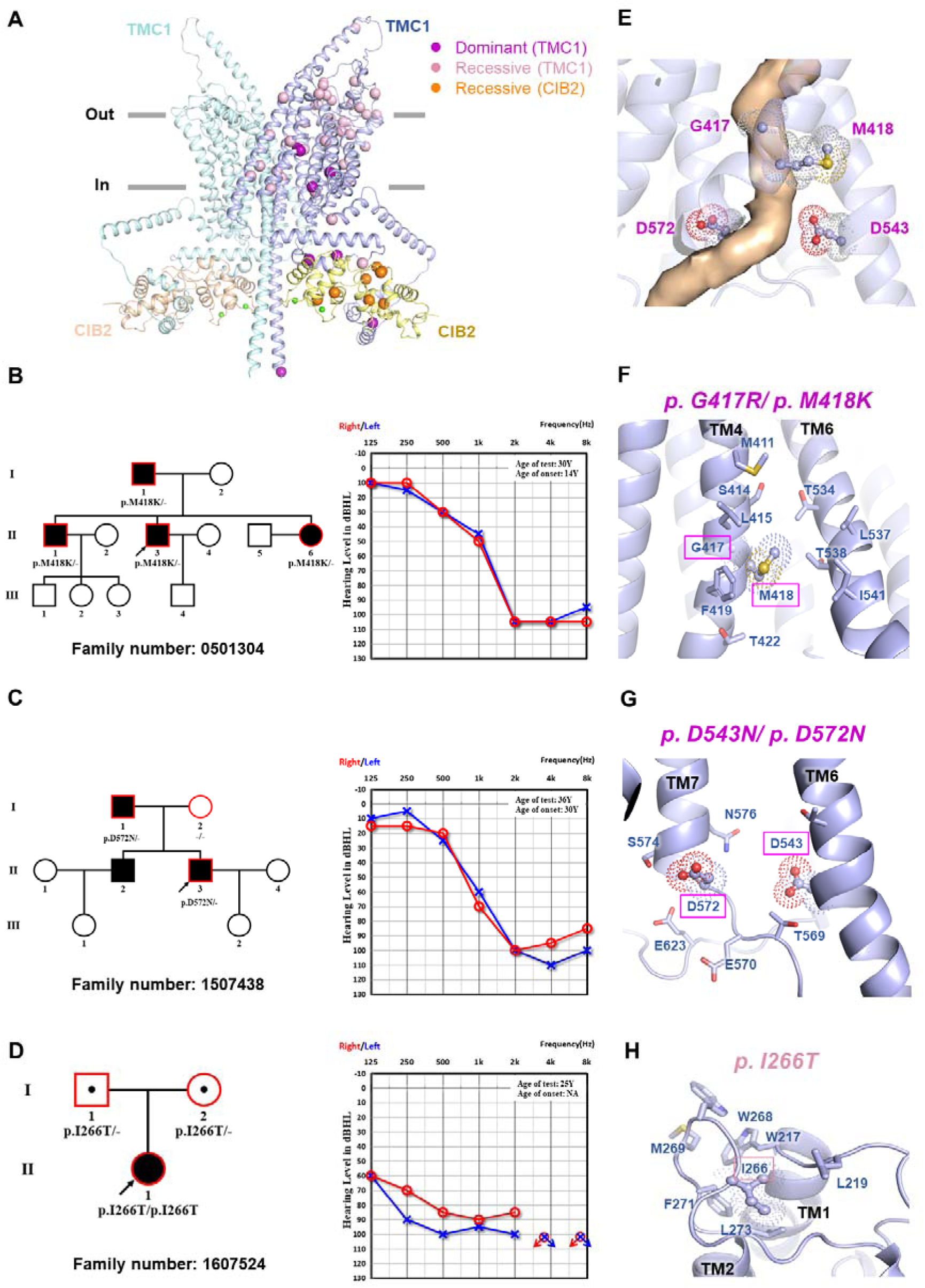
The autosomal dominant hearing loss mutations of TMC1 around the putative pore of the MET. **(A)** The locations of disease mutations were mapped onto the structure model of TMC1-CIB2. **(B, C, D)** Left panel: The pedigrees of families 0501304 and 1507438 with autosomal dominant inheritance pattern (M418K and D572N), and family 1607524 with autosomal recessive inheritance pattern (I266T). The arrows pointed to the probands, and the family members with genetic testing were framed in red; Right panel: Audiological phenotypes of deafness patients with pure tone audiograms of the probands. The red represents the hearing of the right ears, and the blue represents the left ears; symbols “o” and “x” denote air conduction pure tone thresholds at different frequencies in the right and left ears. dB: decibels; Hz: Hertz; Y: years old. (**E**)The structural details of the putative pore of TMC1. (**F, G, H**) Structure analysis on the dominant variants (framed in purple) and recessive variants (framed in pink). Residues involved in protein interaction are shown with a stick model, and variants residues are shown as dots.

This study recruited three hearing loss families to reinforce the evidence of the genetic heterogeneity of *TMC1*. The *TMC1* pathogenic mutations were confirmed and co-segregated in all three families. Family 0501304 (p.M418K) and 1507438 (p.D572N) showed an autosomal dominant inheritance pattern (Fig. 5B&5C), and family 1607524 (p.I266T) showed an autosomal recessive inheritance pattern (Fig. 5D). Patients in family 0501304 (p.M418K) and 1507438 (p.D572N) showed a late-onset bilateral sensorineural severe hearing loss, mainly affecting high-frequencies. The patient in family 1607524 (p.I266T) showed an early-onset bilateral profound sensorineural hearing loss in which all frequencies were affected.

The total of 51 missense variants (43 in TMC1 and 8 in CIB2) were mapped onto the complex structure of mammalian TMC1-CIB2 (Fig. 5A, Table S2&S3). Truncation/deletion mutations are not discussed here as these can be interpreted as null mutations. Among the 43 TMC1 mutation site, 14 are located in the extracellular (EC) region, 24 in the transmembrane (TM) region, and 5 in the cytosolic (Cyto) region (Fig. 5A&Table S3). For the mutants in the EC region, further investigation is needed to determine the involvement of binding to other components of MET.

Interestingly, all of the pore-region mutants are linked to autosomal dominant hearing loss. Four of the 24 TM variants occur in the putative pore of TMC1 (G417R and M418K in TM4, D543N in TM6, and D572N in TM7) (Fig. 5E). These disease-causing mutations in this category are likely to impact the conductivity of the putative pore of TMC1. As shown in Fig. 5F, p. G417 and p. M418 are neighbors in TM4 of TMC1. Substituting Gly417 with Arg and Met418 with Lys is expected to disrupt the hydrophobic networks formed by the surrounding residues and introduce positively charged residues that are unfavorable for Ca^2+^ permeation through the channel pore (Fig. 5F). The other two dominant mutations, p.D543N in TM6 and p.D572N in TM7, are located near the cytosolic end of the putative channel pore. Substituting Asp with Asn would remove the negatively charged residue, which is critical for binding Ca^2+^ (Fig. 5G). Previous studies have shown that equivalent mutations of M418K in mouse (p.M412K) decreases Ca^2+^ permeability and D569N in mouse (equivalent to p.D572N in human) diminished the influx of Ca^2+^ ^39^. Moreover, p.D572N in human TMC1 disrupted LHFPL5 binding and destabilized TMC1 expression^65^. The recessive mutation p.I266T lies at the extracellular loop between TM1 and TM2 and forms extensive hydrophobic interactions with W217, L219, W268, M269, F271, and L273. Substitution of Ile with Thr may influence these hydrophobic interactions (Fig. 5H) and result in a recessive phenotype.

### Hearing loss-related Mutations disturb TMC1-CIB2 interaction

Notably, the remaining dominant mutations (p.M113I, p.F313S, p.S320R, and p.R759H) in TMC1 are in the binding regions with CIB2 (Fig. 6A), suggesting the interaction of TMC1 and CIB2 plays critical roles in MET. As mentioned above (Fig. 1E), M113 (equivalent to M107 in mouse) at CR1 directly participates in binding with CIB2, and its sidechain inserts into the hydrophobic groove formed by Y16, C19, and V92 of CIB2. The groove would not accommodate the substitution of Met with Ile (Fig. 6C). p.F313S and p.S320R (equivalent to p.F307 and p.S314 in mouse), located at the binding interface of CR2 and CIB2, are expected to destabilize the hydrophobic contacts formed by F313, F318, W321, I338, F342, I346 from TMC1 and F70, Y115, L135, V148 from CIB2, thus impairing the interaction of TMC1 CR2 and CIB2 (Fig. 6D). As expected, these mutations all weakened or even abolished the TMC1-CIB2 interaction in ITC-based assays (Fig. 6B&S7D).

**Fig. 6.**
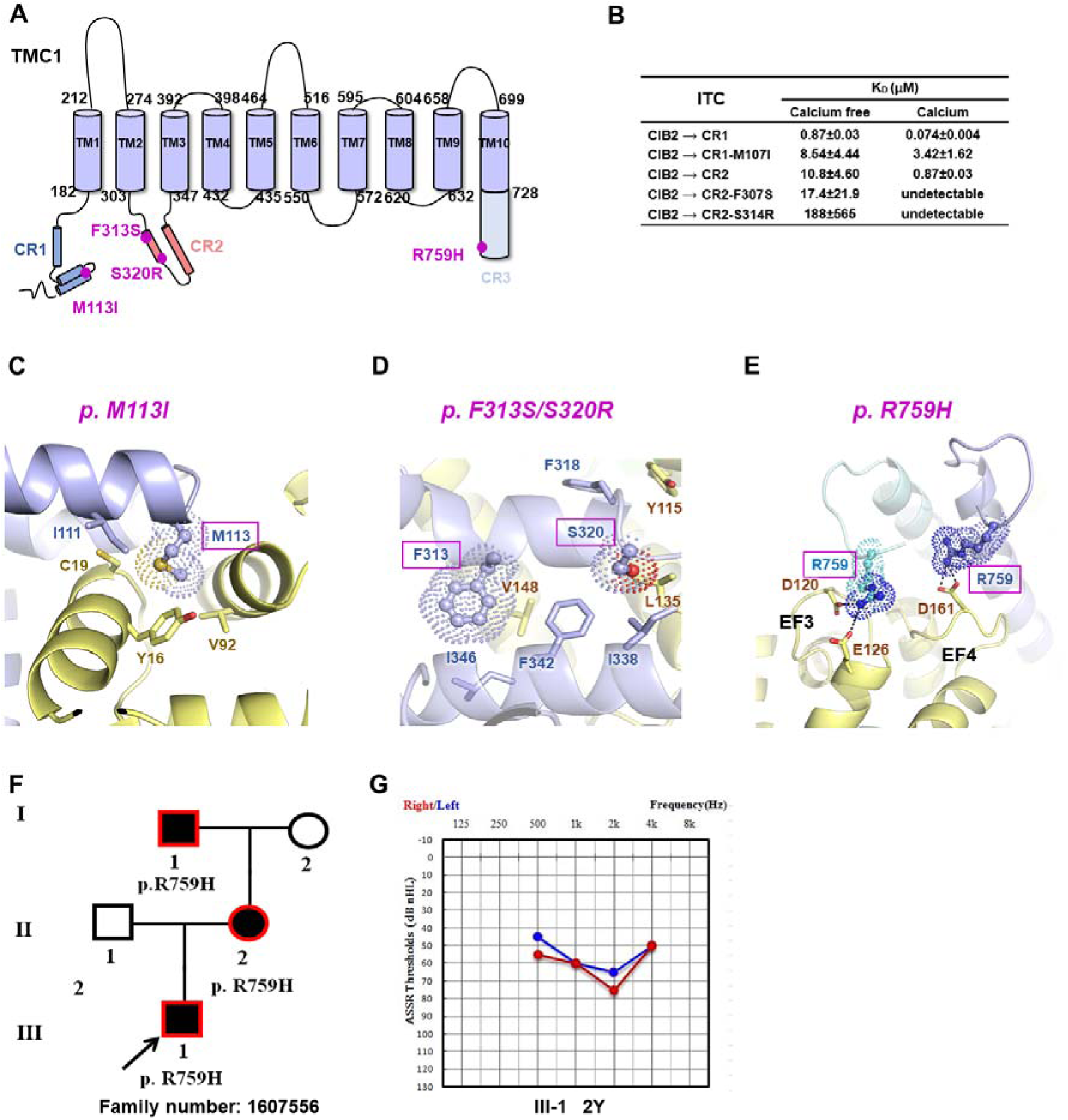
Dominant mutations on the cytosolic regions of TMC1. (**A**) A domain diagram of TMC1 with labeled dominant disease mutations colored in purple. (**B**) A table summarizes the ITC results of CIB2 interacting with variants of TMC1. (**C-E**) The structural details of the dominant variants of TMC1 interacting with CIB2. Dominant variants are framed in purple and shown as dots, and residues involved in protein interaction are shown with a stick model. (**F**) The pedigree of the family 1607556 with autosomal dominant inheritance pattern. The arrows pointed to the probands, and the family members with genetic testing were framed in red. (**G**) Auditory steady-state response (ASSR) hearing threshold of the probands tested at two years old. The red represents the hearing of the right ears, and the blue represents the left ears. dB: decibels; Hz: Hertz; Y: years old.

**Fig. 7.**
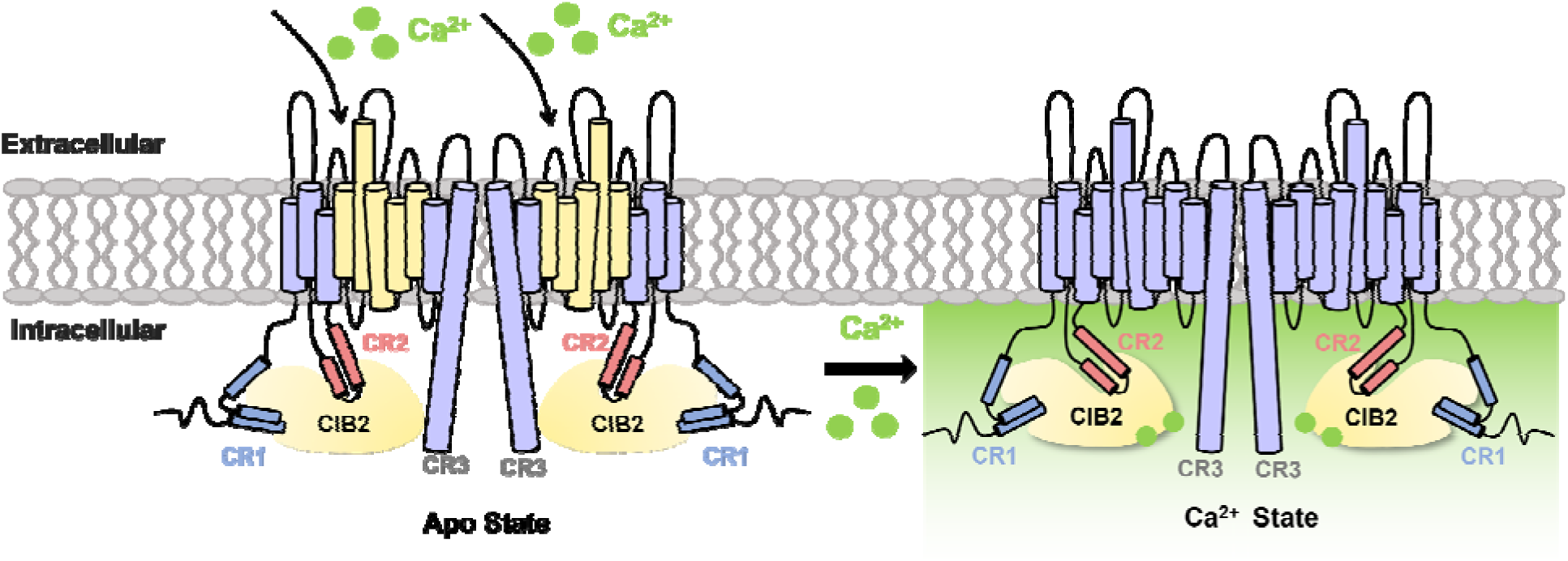
The model of Ca^2+^ mediating the TMC1-CIB2 complex.

The p.R759H missense mutation resides at the C-terminal end of TMC1 CR3. According to the MD-derived model of TMC1-CIB2 *apo* form, the two R759 residues from the TMC1 dimer form salt bridges with the residues for Ca^2+^ coordination in CIB2 (i.e., D120^EF3^, E126^EF3,^ and D161^EF4^) (Fig. 6E). Substitution of Arg759 with His is expected to interfere with the salt bridges’ formation, further destabilizing the binding of CR3 and CIB2. Family 1607556 (p.R759H) indeed showed an autosomal dominant inheritance pattern (Fig. 6F), and the patient showed an early-onset bilateral profound sensorineural hearing loss at two years old, mainly affecting middle-frequencies (Fig. 6G). The discovery of a human mutation on the binding interface of TMC CR3-CIB2 reinforces our model, highlighting the pathophysiological significance of this newly identified CR3 binding site.

Eight deafness mutations in CIB2 have been identified until now (Fig. S6A). To further explore the effects of the mutations on CIB2-TMC1 binding, we picked up four mutations (p.E64D, p.R88W, p.F91S, and p.I123T) and performed ITC assays. Unsurprisingly, all four CIB2 mutations showed decreased binding affinities with TMC1 CR1 or CR3 (Fig. S6F). The structural analysis allows us to rationalize the structural effects of these mutations. E64D missense mutation is located at the h3 helix and forms salt bridges with R33 located at the h2 helix of CIB2, therefore a lack of methyl might weaken the interaction and destabilize the folding of CIB2 (Fig S6B). The F91S missense mutation, located at the h5 helix of CIB2, is expected to interfere with the hydrophobic contacts among L30, F34, L37, M62, F88, F91, V92, and F95, thus destabilizing the folding of CIB2 (Fig. S6C). Similarly, p.I123T near the Ca^2+^ binding site might also impair the surrounding hydrophobic networks (Fig. S6D). The mutation R66W occurred on the hydrophilic loop between h3 and h4 and is close to another deafness mutation site R186W in three-dimensional space (Fig. S6E). Given that the sidechains of Arg88 and Arg186 are exposed to solvent, the substitutions of Arg with the large hydrophobic sidechain of Trp are speculated to destabilize the protein (Fig. S6E). Furthermore, it has been recently reported that R186W mutant significantly slows down MET channel activation and causes loss of membrane-shaping protein BAIAP2L2 localization at the lower tips of stereocilia^66^. Although these mutations are not directly involved in interacting with TMC1, they might impair the overall conformation of CIB2, thus interfering with its binding capacity.

In summary, four dominant variants of TMC1 cluster around the putative ion channel pore, while others are located in the cytosolic regions responsible for binding with CIB2. With the structural and biochemical analyses on the variants and investigation on the pedigrees and audiological phenotypes of deafness patients, we have revealed this genotype - structype - phenotype correlation. This understanding can facilitate accurate diagnosis and tailored treatment options for individuals affected by TMC1-related hearing loss.

## DISCUSSION

Our study reveals that the TMC1-CIB2 complex undergoes a Ca^2+^-induced conformational change, highlighting the crucial role of Ca^2+^ in MET. In addition to the previously identified TMC CR1 binding to CIB2^29^ and the reported complex structure of TMC1 CR2-CIB3^30^, we identified a new binding site on TMC1, named CR3, that only interacts with *apo* CIB2. Notably, the TMC1 CR3 region is conserved only among vertebrates, coinciding with the emergence of the inner ear. Our findings suggest that this newly discovered CR3 binding is also pathophysiological critical, as human mutations on the binding interface are linked to hearing loss (Fig. 6E&6F&6G).

We assembled the intact mammalian TMC1-CIB2 complex, using machine-learning algorithm and our high-resolution TMC1 CR1-CIB2 structure. Based on this model, we developed an algorithmic molecular dynamic-based simulation workflow to examine the molecular dynamic behavior of the TMC1-CIB2 with/without Ca^2+^, revealing that the complex undergoes a Ca^2+^-induced conformational change (Fig. 2A&S3A&7), which was further supported by the in-vivo mouse model (Fig. 4) and clinical genetic data (Fig. 5&6).

### EF-hand in Sensing systems

The EF-hand motif is a common calcium-sensing motif, participating in various physiological processes. The classical EF-hand is a helix-loop-helix motif, with the loop region accommodating calcium. Calmodulin is the most famous EF-hand protein, which serves as the Ca^2+^ sensor for several voltage-gated sodium and calcium channels. At elevated cytosolic Ca^2+^ levels (caused by channel opening), Ca^2+^-bound calmodulin changes its binding mode with the calcium channel and inactivates the channel rapidly. TRP channels generally act as molecular sensors of multiple stimuli, ranging from changes in pH, chemical agents, and temperature. TRPs are unique by including the EF-hand region in its cytosolic sequence instead of interacting with other EF-hand-containing proteins. CIB2, just like calmodulin, has four EF-hands, with two capable of binding calcium. Here, we discovered the Ca^2+^ regulation on the TMC1-CIB2 complex. We propose a model here: in the *apo* state (without Ca^2+^), CIB2 binds to CR1, CR2, and CR3 of TMC1. Following the influx of Ca^2+^, CIB2 dissociates from CR3 and induces an overall conformation change in association with CR1 and CR2, which may be coupled with the closure of the ion pore. Compared to other mechanosensitive channels, the hearing MET channel has ultrafast open-close kinetics for fluctuating sound input. Calcium ion has a high-speed diffusion rate, so it is widely involved in various rapid regulation processes. Our study provides insights into Ca^2+^-mediated gating for sound perception by combining in-vitro, in-vivo, and clinic data.

It is recognized that Ca^2+^ plays a crucial role as a regulator in MET kinetics. The influx of Ca^2+^ can modulate the activation and adaptation of the transducer current in a concentration-dependent manner^44,67^. However, the involvement of calcium in fast adaption of MET remains under debate^68–70^. The discrepancies observed could arise from different stimulation configurations of electrophysiology experiments. Kinetics studies on the Ca^2+^ binding of TMC1-CIB2 will contribute to the topic for discriminating the fast/slow reactions, in regard of the timescale of the transduction occurs (tens or even hundreds of kilohertz). Further research is needed to elucidate the mechanisms underlying the modulation of the MET channel across different time scales.

### Linking Genotype, Structype (structure type), and Clinic Phenotype together

The issue of genetic heterogeneity has long posed a challenge in clinical diagnosis, particularly in non-syndromic hereditary hearing loss, which has a high degree of genetic heterogeneity, especially for *TMC1* with dominant and recessive inheritance patterns. Depending on the positions on the protein structure and the physicochemical properties of the substituted amino acid, variants can affect the proper function of the protein in different ways. For the recessive mutations, the disturbance caused by the amino acid change can be compensated by the wild-type copy. But some variants can be severe and have a dominant negative effect. In that case, one copy of the substation is enough to cause the phenotype as an autosomal dominant disease. Autosomal recessive hearing loss (DFNB7/11) and autosomal dominant (DFNA36) are like the two faces of *TMC1*.

To date, over 100 pathogenic variants in TMC1 have been reported^71,72^, with only 8 of them responsible for DFNA36 (M113I, F313S, S320R, G417R, M418K, D543N, D572N and R759H). However, the mechanism by which the dominant and recessive mutations lead to abnormal TMC1 function and regulation remains mostly unexamined. We collected 43 missense mutations of hearing loss patients from online databases as well as clinic data and mapped these mutation sites on the TMC1-CIB2 complex structure model (Fig. 5A). We systematically characterized the deafness mutations by combining structural analysis, biochemical validation, *in-vivo* assays, and genetic studies, thoroughly investigating the clinical phenotypes of hearing loss from micro to macro (Fig. 5&6&S6). Four of the eight dominant mutation sites cluster around the ion pore, while the other four are responsible for binding with the Ca^2+^ sensor CIB2. This suggests that all the dominant mutants are directly associated with the gating process of the mechanoreception transduction channel for sound perception. This provides an atomic level mechanism for the genetic heterogeneity of *TMC1* in hearing-loss. Linking genotype, structype (structure type), and clinic phenotype together will bring the study of MET in hearing to a new era.

## MATERIALS AND METHODS

### Plasmids Construction, protein expression, and purification

We produced different constructs of CIB2 (reference number UniProt: Q9Z309) and TMC1 (reference number UniProt: Q8R4P5) using standard PCR (Vazyme, Nanjing, China) and homologous recombination (Yeasen, Shanghai, China). All point mutations and disease mutations of CIB2 and TMC1 described in this study were introduced using the PCR-based mutagenesis method. Both primers and DNA sequencing were provided by GENEWIZ (Suzhou, China) and BioSune (Shanghai, China). CIB2, TMC1 CR1 (amino acids 81-130), TMC1 CR2 (amino acids 298-352), TMC1 CR3 (amino acids 718-757), and all point mutations and disease mutations of CIB2 and TMC1 were cloned into the appropriate expression vectors, pET.32m.3c with trx-6xhis tag, pMal-c2X with the mbp-6xhis tag, pEGFP-C1 with eGFP tag or pFLAG-CMV2 with flag tag, as required.

Recombinant proteins were expressed in *Escherichia coli* BL21 (DE3) cells grown in LB medium at 37°C until the OD_600_ values reached ∼0.5. The BL-21 cells were then induced with 1mM IPTG and incubated at 16 for ∼16-20 hours. Recombinant proteins were purified using Ni^2+^-NTA agarose affinity column followed by size-exclusion chromatography (SEC) (Superdex 200 26/600 column from GE Healthcare). The purification buffer contained 50 mM Tris pH 7.8, 100 mM NaCl, 1 mM dithiothreitol (DTT), and 10 mM ethylenediaminetetraacetic acid (EDTA). In calcium state the buffer changes to 50 mM Tris, pH 7.5-8.0, 100 mM NaCl, and 10 mM CaCl2. During crystallization, SEC-MALS, and NMR-based titration experiments, the N-Terminal Trx-His6 tag was removed by 3C protease and another step of SEC chromatography.

### Crystallization, data collection, and structure determination

We did a point mutation at CIB2 p.E141 (mutated to Asparagine) to obtain high quality diffraction data (Fig. S1&S7B, Table. S1). CIB2-E141N and TMC1 CR1 were copurified and were concentrated to a final concentration of 10 mg/ml for crystallization. Initial crystallization screening was performed with 96 conditions kits from HAMPTON. The complex grew at 16°C by the vapor diffusion method in sitting drops, in buffer containing 20% v/v 2-Proranol, 0.1 M MES monohydrate pH 6.0, and 20% w/v PEG monomethyl ether 2000. Crystals were cryoprotected in a reservoir solution with 20% glycerol. Diffraction data were collected at BL19U1 at Shanghai Synchrotron Radiation Facility (SSRF, Shanghai, China) and were processed with HKL3000^73^.

The structure was solved by PHASER in CCP4^74^. The crystal structure of CIB3 (PDB code: 6WU5) was used as the search model. CCP4 Autobuild was used to improve the structure completeness, and further loop building and refinement were performed iteratively using Coot^75^ and REFMAC PHENIX^76^. The final refinement statistics of the structure are listed in Table S2, and structural diagrams were prepared using PyMOL (http://www.pymol.org).

### Isothermal titration calorimetry assay

The ITC measurements were carried out on a MicroCal ITC200 system (Malvern Panalytical) at 25°C. Approximately 300-500 μM of TMC1 truncations were loaded into the syringe and titrated into the cell ∼30-50 μM protein of CIB2 WT or mutations. 10 mM EDTA/CaCl_2_ were added in buffer. The concentration of Ca^2+^ or Mg^2+^ used in Fig S2A was 2 mM; in Fig S2B was 5mM. We constructed TMC1 CR2 with maltose binding protein (MBP) tag to get monomer protein. Each titration point was acquired by injecting a 1-μl aliquot of the protein sample from the syringe into the protein sample in the cell. Titration data was analyzed using MicroCal PEAQ-ITC Analysis Software and fitted by the one-site binding model.

### Inductively coupled plasma-Mass Spectrometry (ICP-MS)

The ICP-MS measurements were carried out on a NexION2000 (Flexar20 HPLC) system at 25.

We prepared CIB2 and CIB2 & TMC1 CR1 complex samples with 50 mM Tris pH 7.8, 100 mM NaCl, 1 mM dithiothreitol (DTT), and 10 mM ethylenediaminetetraacetic acid (EDTA) as described above to eliminate metal ions influences. The Trx-tags were removed by another SEC step using the same buffer without EDTA to remove the excess EDTA. The samples were concentrated to 300 μM, adding 1:1, 1:3, 1:5 or 1:10 molar radio with CaCl_2_ or MgCl_2_. Then we used desalting columns (GE Disposable PD-10) to remove the excess Ca^2+^ or Mg^2+^. The ICP-MS measurements were performed by the Instrumental Analysis Center of SJTU, and data analysis was by GraphPad Prism 7.0.

### SEC coupled with multi-angle static light scattering

A superose 12 10/300 GL column (GE Healthcare), a multi-angle static light scattering detector (miniDAWN, Wyatt), and a differential refractive index detector (Optilab, Wyatt) were coupled with the AKTA FPLC system. The column was pre-equilibrated overnight, and purified proteins were prepared at a concentration of ∼300 μL and 100 μM for loading. Data was analyzed using ASTRA 6 (Wyatt).

### NMR spectroscopy

Nuclear Magnetic Resonance (NMR) samples of CIB2 of 300 μM in complex with TMC-CR1 (residues 81-130) of 300 μM were prepared in buffer containing 50 mM Tris pH 7.0, 100 mM NaCl, 10 mM EDTA (*apo* state) or 10 mM Cacl2 (calcium state) in 10% v/v D_2_O. All NMR spectra were collected at 300K on Agilent VnmrS DD2 700MHz spectrometer equipped with a 5mm XYZ PFG HCN room temperature probe. Chemical shifts were referenced to external DSS. Spectra were processed using the program NMRPipe and analyzed with the program NMRFAM-SPARKY.

### Co-immunoprecipitation and western blotting

HEK 293 cells were maintained at 37 in 5% CO2 using DMEM (HyClone) supplemented with 10% FBS (Gibco), and 1% penicillin–streptomycin (Invitrogen). Cells were transfected with 3μg of each plasmid using Lipo 3000 (Invitrogen). After 24 hours, cells were washed with 1x PBS, then centrifuged at 12000 rpm for 10 min. Cells were lysed using a modified RIPA buffer containing 50 mM Tris (pH 8.0), 150 mM NaCl, 5 mM EDTA (*apo* state) or 5 mM CaCl_2_ (calcium state), 1% Triton, 0.1% SDS and a Roche Complete Protease Inhibitor Tablet. Then cell lysates were rotated for 30 min at 4 followed by centrifuged at 12000 rpm for 10 min at 4. Cell lysates were immunoprecipitated for 1 hour at 4 using 20 μL Anti-GFP Affinity Beads (Smart-Lifesciences, Changzhou, China). After immunoprecipitation, cell lysates were centrifuged at 3000 rpm for 3 min at 4 and washed three times with RIPA buffer. 2x Loading Buffer was added into lysates and boiled for 10 min. SDS-PAGE was performed using 4-20% Tris-Glycine gel (Beyotime, Shanghai, China) and then transferred to PVDF membranes (Millipore). Membranes were block for 2 hours with 5% non-fat dry milk in 1x PBST containing 25 mM Tris (pH 7.5), 150 NaCl and 0.1% Tween-20. Membranes were incubated with Primary antibodies at 4 overnight and incubated with secondary antibodies at RT for 1 hour after washed three times with 1x PBST. Membranes were finally washed and imaged with Chemiluminescent HRP Substrate (Millipore).

Primary antibodies were as follows: mouse anti-GFP (1:5000, Santa Cruz), rabbit anti-FLAG (1:4000, Cell signaling), mouse anti-GAPDH (1:5000, Proteintech, Wuhan, China). Secondary antibodies were goat anti-mouse (1:5000, Invitrogen) and goat anti-rabbit (1:5000, Invitrogen).

### Injectorporation and whole-mount immunostaining

Injectorporation and whole-mount immunostaining was performed as previously described^77^. Briefly, cochlear sensory epithelia were dissected out of P3 wild-type mice followed by culturing in DMEM/F12 with 1.5 μg/mL ampicillin. GFP-CIB2 expression plasmids (1 μg/μL in 1× Hanks’ balanced salt solution) were delivered to the sensory epithelia using a glass pipette (2 μm tip diameter). A series of three pulses at 60 V lasting 15 ms at 1-s intervals were applied to cochlear tissues using an electroporator (ECM Gemini X2, BTX, CA). After culturing for 1 day *in vitro*, the samples were fixed with 4% paraformaldehyde (PFA) in PBS for 20 min, then permeabilized and blocked with PBST (1% Triton X-100, and 5% donkey serum in PBS) for 1 hour. To visualize the stereociliary F-actin core, samples were then incubated with TRITC-conjugated phalloidin (Sigma-Aldrich, Cat. No. P1951, 1:1000 dilution) in PBS for 20 min at room temperature. The samples were mounted in PBS/glycerol (1:1) and imaged with a confocal microscope (LSM 900; Zeiss).

### Whole-cell electrophysiology of hair cells

Cochleae of P3 *Cib2^-/-^* mice were dissected in dissection solution, which contains 141.7 mM NaCl, 5.36 mM KCl, 0.1 mM CaCl_2_, 1 mM MgCl_2_, 0.5 mM MgSO_4_, 3.4 mM L-glutamine, 10 mM glucose, and 10 mM H-HEPES (pH=7.4). After injectoporation as described above, the basilar membrane was transferred into a recording chamber with recording solution, which contains 144 mM NaCl, 0.7 mM NaH_2_PO_4_, 5.8 mM KCl, 1.3 mM CaCl_2_, 0.9 mM MgCl_2_, 5.6 mM glucose, and 10 mM H-HEPES (pH=7.4). These acutely isolated sample need be recorded within 1 hour. Recording was performed under an upright microscope (BX51WI, Olympus, Tokyo, Japan) with a 60 × water immersion objective and an sCMOS camera (ORCA Flash4.0, Hamamatsu, Hamamatsu City, Japan) controlled by MicroManager 1.6 software. Patch pipettes resistances need to be controlled within 4-6 M’Ω. Intracellular solution contained (in mM): 140 mM KCl, 1mM MgCl_2_, 0.1 mM EGTA, 2 mM Mg-ATP, 0.3 mM Na-GTP, and 10 mM H-HEPES, pH=7.2). Hair cells were recorded with a patch-clamp amplifier (EPC 10 USB and Patchmaster software, HEKA Elektronik, Lambrecht/Pfalz, Germany). MET was recorded by a 10 Hz sinusoidal wave delivered by a 27-mm-diameter piezoelectric disc driven by a home-made piezo amplifier pipette with a tip diameter of 3–5 μm positioned 5–10 μm from the hair bundle. Data were analyzed through Igor Pro 9, with self-made macros.

## ACKNOWLEDGMENTS

We thank staff members of the BL19U1 beamline of the National Facility for Protein Science Shanghai (NFPS) at Shanghai Synchrotron Radiation Facility and Shijing Huang for X-ray diffraction data collection. We thank the staff members of the Large-scale Protein Preparation System and the nuclear magnetic resonance system of NFPS, Shanghai Advanced Research Institute, Chinese Academy of Sciences, China, for providing technical support and assistance in data collection and analysis. We thank Yuyu Guo and Haiyan Yu from the core facilities for life and environmental sciences, Shandong University for the technical support in confocal microscopy. We are also grateful for the support from HPC Platform of ShanghaiTech University.

## Author contributions

S. Wu performed biochemical and structural experiments. L. Lin solved, refined, and analyzed the crystal structures. Q. Hu built the structure model and performed MD simulations. X. Yao, L. Lu and Y. Xi conducted the Immunostaining assays. S. Liu and Q. Liu conducted the MET rescue experiments. H. Wang and Q. Wang collected and analyzed the patient data. S. Wu and W. Zhan performed NMR spectroscopy and analyzed the data. S. Wu, L. Lin, Q. Hu, H. Wang, Z. Xu, F. Bai, and Q. Lu analyzed the data and drafted the paper. All authors commented on the paper. Q. Lu coordinated the project.

## Data and materials availability

The atomic coordinate of the complex of TMC1 CR1 and CIB2 has been deposited to the Protein Data Bank under the accession codes 8Z3F.

